# Elevation of FAM129A in neutrophils exposed to serum of patients with severe sepsis: in silico investigations during a hands on training workshop and follow on validation of protein expression in neutrophils

**DOI:** 10.1101/529446

**Authors:** Jessica Roelands, Laurent Chiche, Radu Marches, Mohammed Toufiq, Basirudeen Ahamed Kabeer, Mohamed Alkhair Ibrahim Alfaki, Marwa Saadaoui, Arun Prasath Lakshmanan, Dhinoth Kumar Bangarusamy, Selvasankar Murugesan, Davide Bedognetti, Wouter Hendrickx, Souhaila Al Khodor, Annalisa Terranegra, Jacques Banchereau, Mathieu Garand, Damien Chaussabel, Darawan Rinchai

**Affiliations:** Sidra Medicine, Doha, Qatar; Hopital Europeen, Marseille, France; Jackson Laboratory for Genomic Medicine, Farmington, CT, USA

## Abstract

Steps involved in reductionist investigation approaches can be imitated using public transcriptome datasets as source of training material. In the present report trainees explored an apparent gap in biological knowledge for FAM129A (family with sequence similarity 129 member A). Elevated abundance of FAM129A transcripts were observed in a transcriptome dataset where neutrophils were exposed in vitro to plasma of patients with sepsis. However, no literature linking FAM129A and either neutrophils, sepsis or inflammation could be identified. Additional datasets were selected to independently validate this initial observation and further explore differential expression of FAM129A in the context of sepsis studies. Follow on investigations carried out at the bench confirmed restriction of the expression of FAM129A protein at the surface of circulating blood neutrophils and monocytes. A potential role for FAM129A in neutrophil survival was inferred from profiling of literature associated with FAM129A, which remains to be investigated in further follow on investigations.

## INTRODUCTION

Sepsis is a life-threatening condition characterized by multiple organ dysfunction [1]. Understanding of the underlying pathogenic mechanism has greatly improved over the past several years. Yet interventions aiming at modulating the exaggerated immune response to infection, one of the root causes of sepsis complications, are yet to materialize in clinical practice. Neutrophils constitute a first line of defense against bacterial infections. This specialized population of leukocytes carries effector functions via the production of immune mediators as well as through microbicidal activity, either resulting from degranulation or through a process that is referred to as “netosis” which consists in formation of extracellular traps that serve to immobilize and kill invading pathogens [2].

In earlier work we have demonstrated the singular ability of neutrophils to respond to immunomodulatory factors present in the plasma of patients with severe sepsis ([3] / GSE49758). This transcriptome dataset which at the time was deposited in the NCBI’s gene expression omnibus was employed more recently in the context of a training workshop conducted at Sidra Medicine in Doha Qatar in November of 2018. Trainees applied reductionist investigation approaches, using available public omics data as a starting point (COD1 training module, as described in: [4]). Individual reports from trainees are included with this submission as supplementary files. Potential gaps in biomedical knowledge were assessed as a first step. It was thus determined that expression of FAM129A by neutrophils had not been reported in the biomedical literature and neither had its induction under inflammatory conditions or in the context of sepsis. The next step consisted in confirming the initial observation in independent datasets and inferring from the literature a potential role for FAM129A in the context of sepsis, inflammation and/or neutrophil immunobiology. Experiments were in addition performed to confirm expression of FAM129A protein in neutrophils.

## MATERIALS AND METHODS

### Workshop format

Briefly, a workshop was organized in November of 2018 at Sidra Medicine in Doha, Qatar. It consisted in applying reductionist investigation approaches while using public transcriptome data as source of training material (COD1 training module [4]). Trainees who joined the workshop carried out hands on activities focusing on two candidate genes, ANXA3 and FAM129A. For each candidate gene activities were guided by two experienced instructors. Manuscript preparation was the primary responsibility of the instructors and of the senior investigator leading the program. Workshop participants prepared each a report which are made available as part of this publication as supplementary files. More details regarding the COD1 training module, including educational material shared during this workshop, will be made available via a separate publication.

### Access to reference datasets

The primary dataset employed as a starting point for identification of gene candidates was deposited in GEO by Khaenam et al ([3], accession GSE49758). This and other reference public transcriptome datasets relevant to sepsis, neutrophil immunobiology or inflammation were deposited in a data browsing application (GXB: http://sepsis.gxbsidra.org/dm3/geneBrowser/list). A total of 48 datasets were included in this curated collection. Each dataset can be individually accessed and profiles of any transcript measure on the array plotted interactively. The GXB tool has been described in details in an earlier publication [5]. Constitution of curated dataset collections is an activity that is undertaken as part of the COD2 training module, as described in a recent review [4].

## RESULTS

### FAM129A levels are increased in neutrophils exposed to septic serum

The “primary” dataset used as a basis for identification of candidate genes was generated and deposited in GEO by our team in the context of an earlier publication [3]. In this experiment neutrophils were isolated from the blood of healthy individuals and exposed in vitro for 6 hours to plasma from adult patients with sepsis. The neutrophil transcriptional response was measured using Illumina beadarrays. FAM129A was prioritized as a target for this training activity on the basis of: 1) increased abundance of this transcripts following in vitro exposure of neutrophils to serum of septic patients (GSE49758, [3]), and 2) the extent of the overlap between the FAM129A literature and literature on sepsis, neutrophil or inflammation. Figure 1 shows levels of FAM129A transcript abundance in neutrophils exposed to plasma from subjects with non-severe or severe sepsis. Difference between these two groups was significant (p <0.01).

**FIGURE 1.**
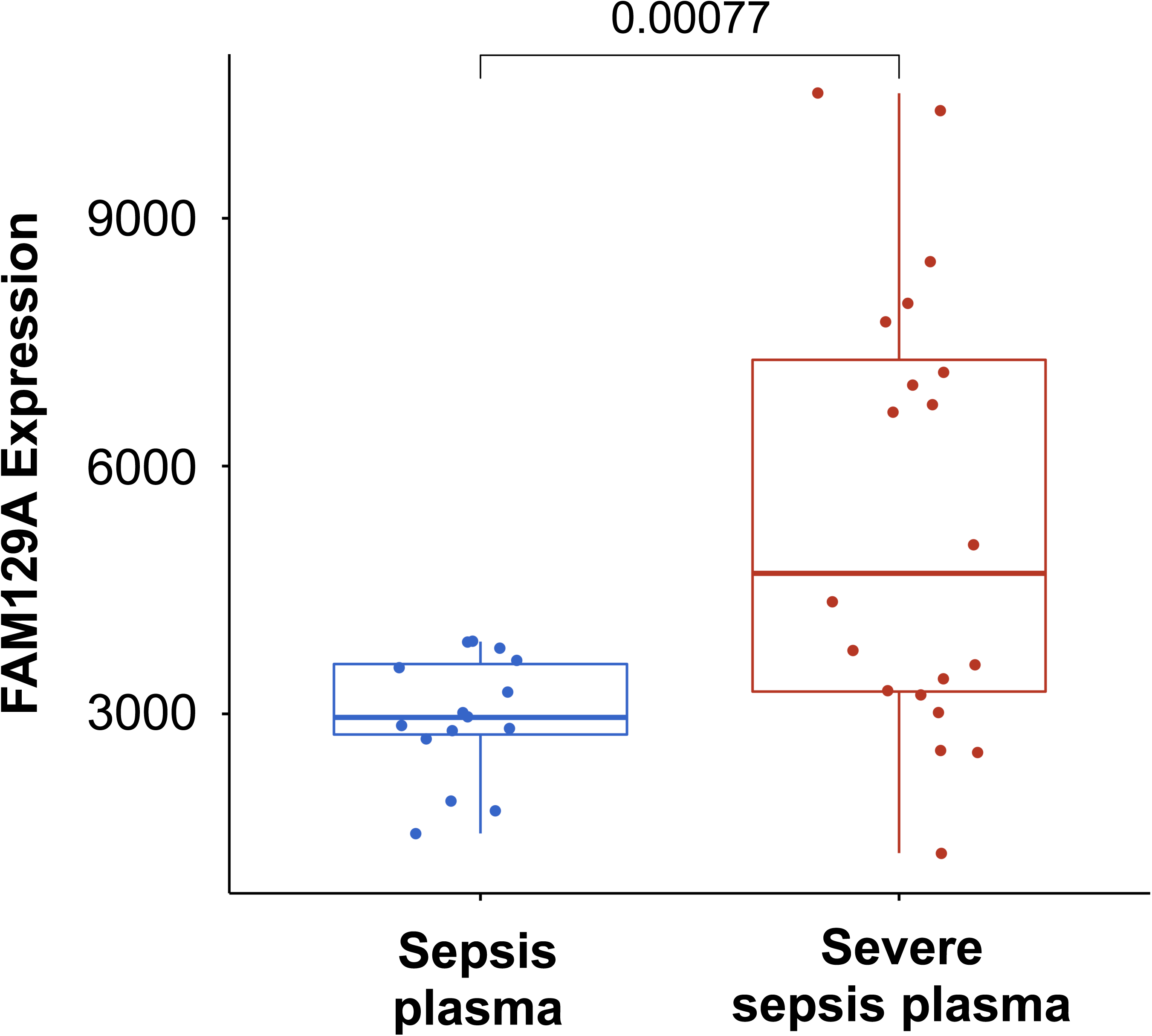
Abundance of FAM129A transcripts is increased in neutrophils exposed to plasma of severe septic patients. Abundance levels of FAM129A extracted from a dataset by Khaenam et al. with accession GSE49755 are plotted on this figure. It shows a significant increase in FAM129A transcript abundance in neutrophils isolated from healthy subjects and cultured for 6 hours in the presence of plasma from patients with severe sepsis, as compared to levels measured in patients with non-severe sepsis.

### Knowledge gap assessment

The next assignment consisted in assessing the extent of the current biomedical knowledge regarding the role of FAM129A in the context of sepsis, inflammation or neutrophil immunobiology. First, a PubMed query was designed that would retreive the FAM129A literature. Official symbol, name and aliases available from the NCBI Entrez Gene database were combined in the following query:

“sequence similarity 129 member A” [tw] OR FAM129A [tw] OR C1orf24 [tw] OR NIBAN [tw]

As of January of 2019, a list of 35 articles was returned when running this PubMed query (listed in **Supplementary file 1**). Next, links between the FAM129A and inflammation, neutrophil or sepsis literature were investigated. The Boolean operator AND was added to the FAM129A query followed by the keywords “Inflammation”, “Neutrophils” or “Sepsis” (Figure 2). No articles among the FAM129A literature intersected with either the neutrophil or sepsis literature. But one of the 35 articles constituting the FAM129A literature intersected with the inflammation literature [6]. In this study FAM129A was identified as one of four genes associated with airway hyper-responsiveness and differentially expressed in laser microdisected airway smooth muscle cells of asthmatic patients vs non-asthmatic controls. The work did not associate FAM129A with neutrophils or sepsis and did not elaborate on the role of FAM129A in the context of atopy or asthma.

**FIGURE 2.**
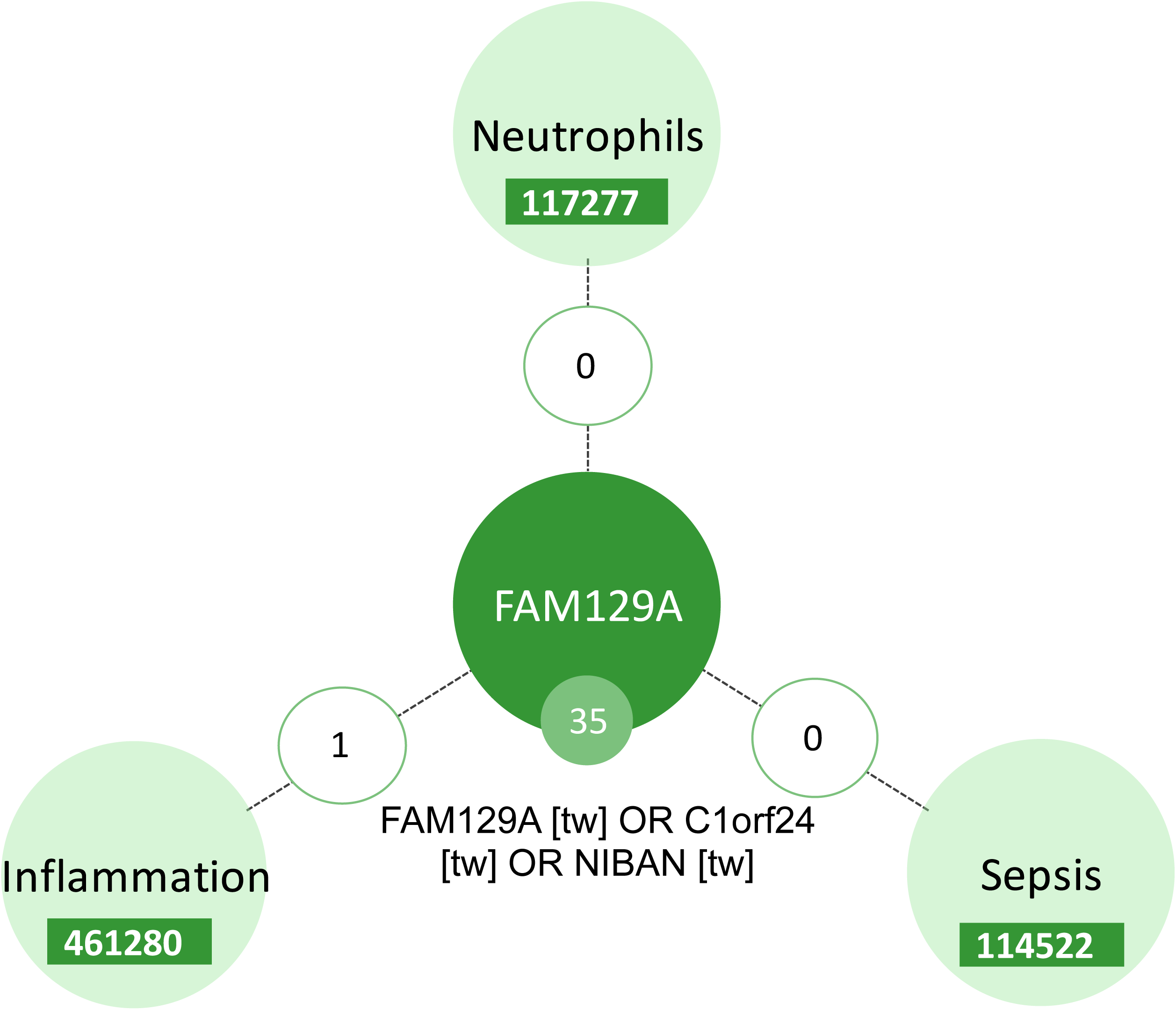
Assessment of gap in FAM129A biomedical knowledge relating to sepsis, inflammation or neutrophil immunobiology. This diagrammatic representation indicates the number of PubMed articles found to be associated with either FAM129A, Neutrophils, Inflammation or Sepsis (each “node”, with numbers of articles provided). Numbers provided on edges connecting each of the notes indicate the extent of the overlap.

### Independent validation in silico

The third task assigned to the trainees was to validate the initial finding in independent public datasets. GEO datasets relevant to sepsis or neutrophil immunobiology were curated and uploaded on the custom GXB data browsing application. This resource comprises 48 datasets and is available via this link: http://sepsis.gxbsidra.org/dm3/landing.gsp. This GXB instance was created expressly in support of the activities conducted in the context of workshops utilizing the same primary dataset and pool of candidate genes (as described in a manuscript that is in preparation, GXB has been previously described and is available on GitHub [5]). Trainees were asked to: first select vdatasets that they estimated to be adequate for validation, on the basis of the information about study/experimental design available in GXB or in the linked primary publication, and, secondly, to access the profile of FAM129A in each of the dataset. This two-step process was established to avoid biasing the validation process. In the case of FAM129A, seven datasets were first selected in which abundance levels of transcript were measured in whole blood, PBMCs or neutrophils of patients with sepsis (Table 1) [7-13]. FAM129A transcript abundance was not measured in one of the datasets [13]. Increase in FAM129A transcript levels in septic patient as compared to controls was found in four of the six remaining datasets (Figure 3). The characteristics of the study populations varied, with three out of six consisting of pediatric subjects and the other three of adults (Figure 4). One study was conducted in Asia, one in Europe and four in North America. Assay characteristics were also different since in four cases samples were run on Illumina beadchips and in two on Affymetrix GeneChips. The datasets in which no increases in levels of FAM129A were not observed were conducted both in adults and in North America. One of them profiled whole blood and used Illumina arrays, while the other profiled neutrophils and used Affymetrix arrays. In yet another dataset in which profiles were generated via RNA sequencing increase in abundance of FAM129A could be observed in whole blood of septic patients as well as in isolated monocytes but it was not the case in neutrophils in which abundance levels were uniformly higher (http://sepsis.gxbsidra.org/dm3/miniURL/view/OF) [14] / GSE60424.

**Table 1:** Validation datasets used to assess levels of FAM129A RNA abundance in septic patients.

| GSE ID | Title | Age | Array platform | Population geographic location | Sample type | Group A | Group B | Exp A | Exp B | B/A | t-test | F-test |
| --- | --- | --- | --- | --- | --- | --- | --- | --- | --- | --- | --- | --- |
| **GSE13015** | Genomic Transcriptional Profiling Identifies a Blood Biomarker Signature for the Diagnosis of Septicemic Melioidosis-GSE13015-Healthy-Melioidosis-Other Sepsis (39) – PMID: 19903332 | Adult | Illumina | Asia | Blood | Healthy & T2D | Melioidosis & other sepsis | 3149 | 8284 | 11.4 | <0.001 | <0.001 |
| **GSE16129** | Enhanced Monocyte Response & Decreased Central Memory T Cells in Children with Invasive Staphylococcus aureus Infections – PMID: 19424507 | Pediatric | Affymetrix | North America | PBMCs | Healthy | S. aureus | 315 | 1170 | 3.7 | <0.001 | <0.001 |
| **GSE25504** | Whole blood mRNA expression profiling of host molecular networks in neonatal sepsis (63) – PMID: 25120092 | Pediatric | Illumina | Europe & Africa | Blood | Control | Neonatal sepsis | 729 | 1718 | 2.4 | <0.001 | <0.001 |
| **GSE30119** | Genome-wide analysis of whole blood transcriptional response to community-acquired Staphylococcus aureus infection in vivo (39) PMID: 22496797 | Pediatric | Illumina | North America | Blood | Healthy | S. aureus infection | 2553 | 4050 | 1.6 | <0.001 | <0.001 |
| **GSE64457** | Marked alterations of neutrophil functions during sepsis-induced immunosuppression (23). PMID: 26224052 | Adult | Affymetrix | Europe | Neutrophils | Healthy control | Septic Patient | 7512 | 6880 | 0.9 | 0.17 | 0.02 |
| **GSE54514** | Whole blood transcriptome of survivors and non survivors of sepsis (163). PMID: 23807251 | Adult | Illumina | Australia/Oceania | Blood | Sepsis survivor and Healthy | Sepsis (Non survivor) | 4655 | 3294 | 0.7 | <0.001 | 0.65 |
| **GSE26378** | Expression data from validation cohort of children with septic shock (103). PMID: 21738952 | Pediatric | Affymetrix | North America | Blood | Normal control | Septic shock patient | N/A | N/A | N/A | N/A | N/A |

**FIGURE 3:**
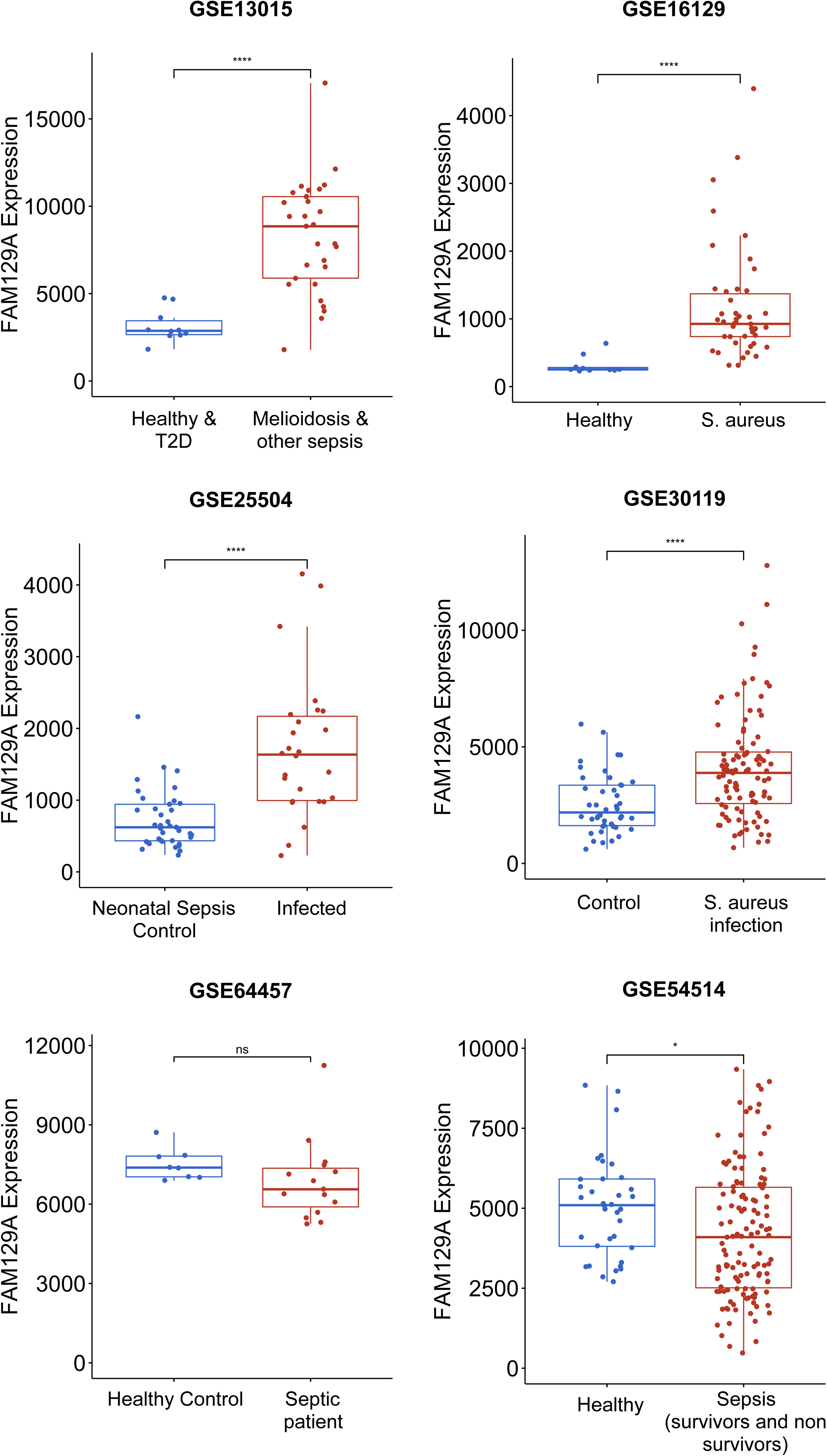
Changes in abundance of FAM129A transcripts in septic patients in six independent datasets. Box plots show levels of FAM129A measured via microarrays in 6 public datasets. Datasets are arranged in the same order as in Figure 2. In each dataset expression levels between septic patients and control subjects are compared.

**FIGURE 4:**
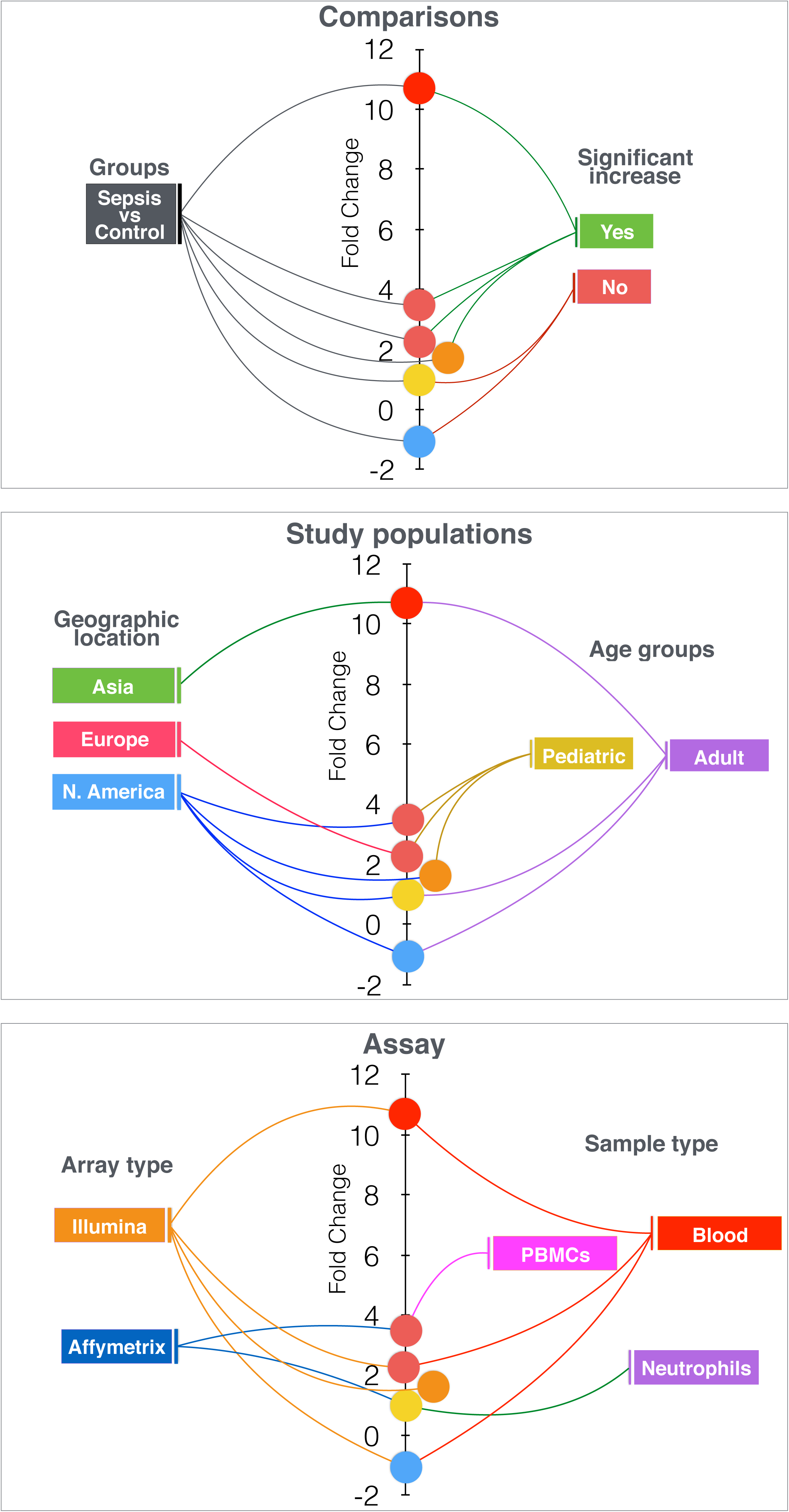
Increase of FAM129A expression in sepsis is observed in four out of six independent datasets. This graph shows fold changes in abundance of FAM129A in six independent datasets selected for follow on investigation of the increase in abundance of FAM129A RNA observed in neutrophils exposed to septic plasma (Figure 1). Each dataset is represented by a spot. The position of the spot on the vertical axis indicates the corresponding fold change in abundance of FAM129A for the dataset. Spots are connected to dataset attributes (metadata). On the top panel attributes relevant to the group comparisons that were performed are shown (groups being compared, whether increase in FAM129A abundance is statistically significant). On the middle and bottom panels attributes describing the study populations and assay, respectively, are overlaid on the same graph. For reference, GEO IDs for the datasets shown on the graph from top to bottom are: GSE13015, GSE16129, GSE25504, GSE30119, GSE64457, GSE54514 (also listed in Table 1). FAM129A transcript abundance profiles for each of these datasets is plotted in Figure 3.

### FAM129A protein is expressed at the surface of monocytes and neutrophils

Further validation results were obtained via flow cytometry analysis of FAM129A protein abundance at the surface of blood leukocyte populations (Figure 5). Whole blood samples were stained with cell-specific markers CD3, CD4, CD8, CD11B, CD14 and CD19. The gating strategy and expression levels of FAM129A relative to an isotype control are shown. These results show highest levels of expression of FAM129A in neutrophils. Uniformly high levels of expression were also observed in monocytes. Levels of expression in B-cell were low, and were the lowest in CD4 and CD8 T-cels.

**FIGURE 5:**
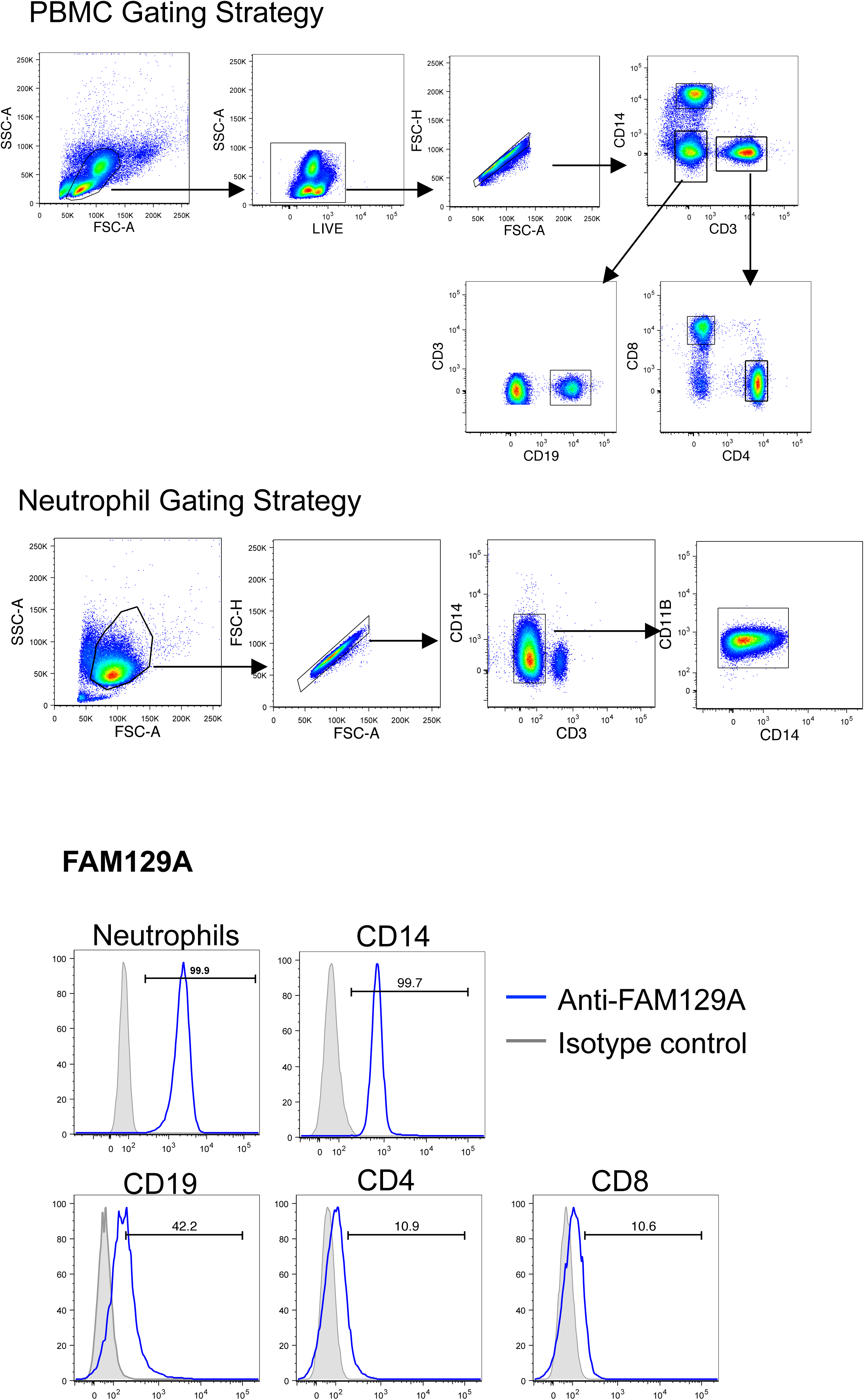
Flow cytometry profiling of FAM129A protein expression on blood leukocyte populations. The top panel describes the gating strategy, which employed a combination of forward and side scatter measurements along with cell surface markers CD3, CD14, CD4, CD8, CD11B and CD19 in order to identify peripheral blood mononuclear cell and neutrophil populations in whole blood. The bottom panel shows levels of expression of FAM129A in neutrophils, monocytes (CD14), B-cells (CD19) as well as CD4 and CD8 T-cells.

### Inferring the role of FAM129A in sepsis and neutrophil immunobiology

The last COD1 assignment requires trainees to build inferences about the potential role for the candidate gene in the context of the primary dataset/study in which the initial observation was made. In order to support this process, the FAM129A literature was profiled in order to identify the main functional “themes” associated with this molecule. Keywords in titles in which FAM129A is mentioned were extracted, categorized and recorded in a spreadsheet. Categories included: “biological processes”, “disease”, “tissue”, “biomolecules”, etc… Subsequently, the prevalence of those keywords in the FAM129A literature was determined. Top keyword or phrases included “thyroid carcinoma” (10 articles out of the 35 constituting the FAM129A literature), which after consolidation with related keyword/phrases such as “thyroid cancer”, “follicular thyroid neoplasms”, “thyroid follicular tumors”, “thyroid tumors” yielded a total of 12 entries. Other major concepts associated with FAM129A were “apoptosis” (7 entries), “proliferation” (6 entries) and “renal tumors” OR “renal carcinogenesis” (5 entries).

Next, these top FAM129A-associated concepts were employed in order to establish indirect relationships with the context of the primary study in which differential FAM129A expression was observed. While only one article could be found connecting FAM129A directly to either the sepsis, inflammation or neutrophil literature, many were found which permitted to establish an indirect connection through via “thyroid cancer”, apoptosis or proliferation (Figure 6).

**FIGURE 6:**
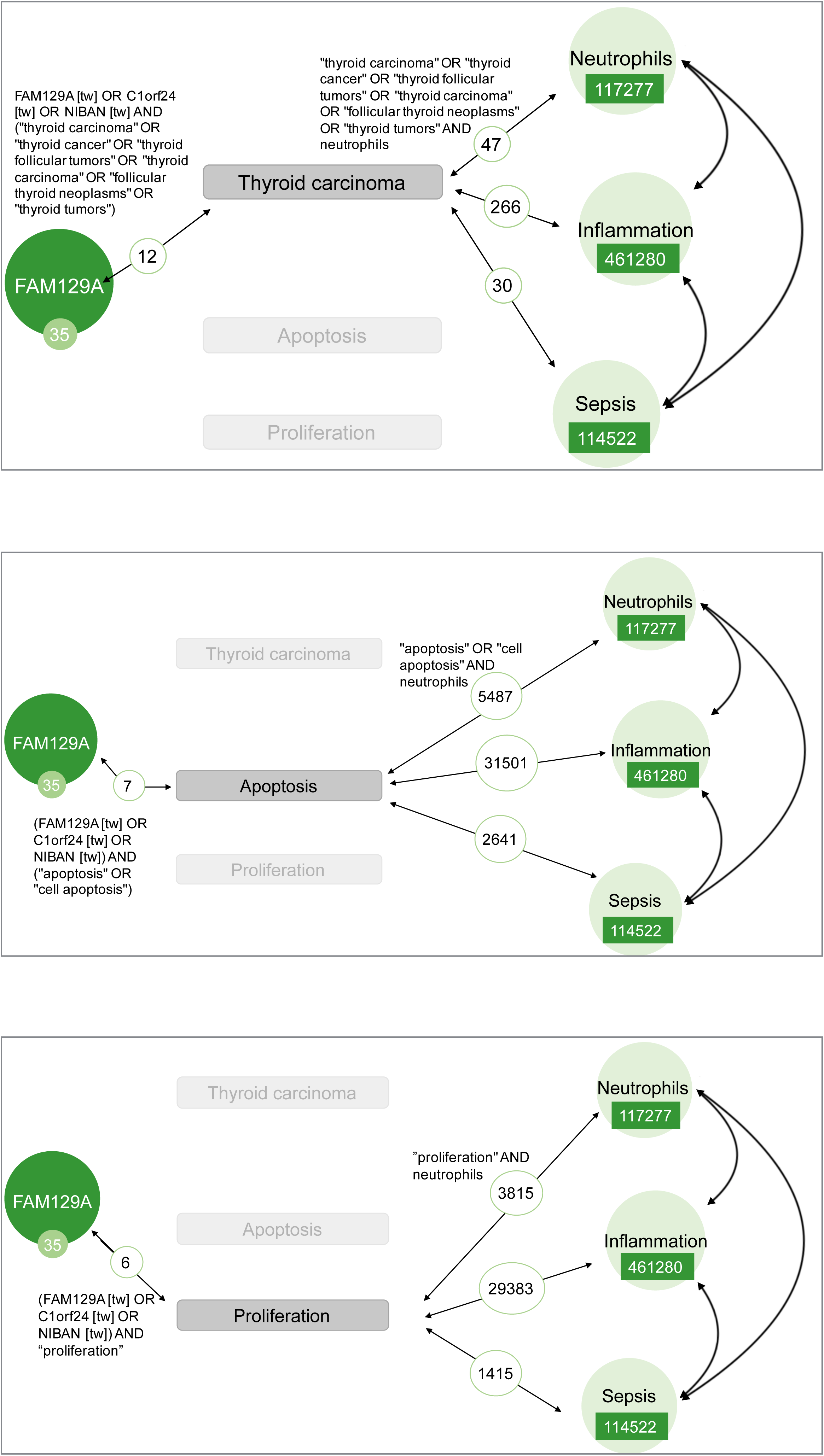
Identification of concepts indirectly linking the FAM129A literature and literature associated with Neutrophils, Inflammation or Sepsis. Each panel indicates the extent of indirect associations established through one of the top concepts associated with the FAM129A literature. In the top panel this concept is “Thyroid carcinoma”. The Pubmed query retrieving the 12 articles in the FAM129A literature in which thyroid carcinoma is mentioned is shown on top of the left most circle. In turn, the query retrieving 47 articles of the neutrophil literature in which thyroid carcinoma is mentioned is shown on the right of this panel. Similar queries were run to retrieve inflammation or sepsis literature mentioning thyroid carcinoma. The middle and bottom panels are organized similarly but explore respectively apoptosis and inflammation as intermediate concepts.

The indirect associations established between FAM129A and sepsis, inflammation and the neutrophil literature were next used as a framework to make inferences about the role of FAM129A in this context as is outlined in the discussion.

## DISCUSSION

While no literature describes the role of FAM129A in neutrophils and its relation to sepsis, the biological concepts apoptosis and proliferation have previously been linked to FAM129A. Expression of FAM129A has been frequently reported in human cancer, including thyroid carcinoma, head and neck cancer and renal carcinoma [15-23]. In human cancer cells, an anti-apoptotic role for FAM129A has been described [16, 24]. A similar role for FAM129A could be possible in neutrophils. Neutrophils have a short life span both in the circulation (8-20 hours) and in tissue (1-4 days). The constitutive neutrophil death is a type of apoptotic death and its regulation is essential for neutrophil homeostasis [25]. A prolonged lifespan of neutrophils has been linked to sepsis in patients with burns, traumatic injuries and pneumonia [26-29]. This could be an explanation for the observed upregulation of FAM129A in neutrophils in association with sepsis or increased severity of sepsis. Thus our hypothesis is that the inhibition of neutrophil apoptosis mediated by FAM129A would lead to an extended life span of neutrophils, subsequently causing damage to healthy tissues. Follow on investigations that could permit testing of this hypothesis include the correlation of FAM129A with p53 expression in neutrophils or assessing FAM129A expression in non-apoptotic and apoptotic neutrophils.

## ACKNOWLEDGEMENTS

Sidra Medicine is a member of the Qatar Foundation for Education, Science and Community Development. The work was supported in part via a grant from the Qatar National Research Fund: QNRF NPRP10-0205-170348

## AUTHOR CONTRIBUTION

Conceptualization: JR, DC, DR. Data curation and validation: DR and MT. Visualization: JR, DC, DR. Analysis and interpretation: JR, LC, RM, MT, BAK, MAIA, MS, AL, DKB, MG, DC, DR. Writing of the first draft: JR, DC, DR. Funding acquisition: DB, DC. Methodology development: MT, BAK, MAIA, MG, DC, DR. Software development and database maintenance: MT. Writing – review & editing: JR, LC, RM, MT, BAK, MAIA, MS, AL, DKB, DB, WH, SAK, AT, JB, MG, DC, DR. The contributor’s roles listed above follow the Contributor Roles Taxonomy (CRediT) managed by The Consortia Advancing Standards in Research Administration Information (CASRAI) (https://casrai.org/credit/).

