## Supplementary file1 for "Elevation of FAM129A in neutrophils exposed to serum of patients with severe sepsis: in silico investigations during a hands on training workshop and follow on validation of protein expression in neutrophils"

Literature returned on January 21, 2019 running the following PubMed query:

"sequence similarity 129 member A" [tw] OR FAM129A [tw] OR C1orf24 [tw] OR NIBAN [tw]

1: Nozima BH, Mendes TB, Pereira GJDS, Araldi RP, Iwamura ESM, Smaili SS,

Carvalheira GMG, Cerutti JM. FAM129A regulates autophagy in thyroid carcinomas in

an oncogene-dependent manner. Endocr Relat Cancer. 2019 Jan 1;26(1):227-238. doi:

10.1530/ERC-17-0530. PubMed PMID: 30400008.

2: Zhu N, Hou J, Wu Y, Liu J, Li G, Zhao W, Ma G, Chen B, Song Y. Integrated

analysis of a competing endogenous RNA network reveals key lncRNAs as potential

prognostic biomarkers for human bladder cancer. Medicine (Baltimore). 2018

Aug;97(35):e11887. doi: 10.1097/MD.0000000000011887. PubMed PMID: 30170380.

3: Evstafieva AG, Kovaleva IE, Shoshinova MS, Budanov AV, Chumakov PM.

Implication of KRT16, FAM129A and HKDC1 genes as ATF4 regulated components of the

integrated stress response. PLoS One. 2018 Feb 8;13(2):e0191107. doi:

10.1371/journal.pone.0191107. eCollection 2018. PubMed PMID: 29420561; PubMed

Central PMCID: PMC5805170.

4: McGeachie MJ, Clemmer GL, Hayete B, Xing H, Runge K, Wu AC, Jiang X, Lu Q,

Church B, Khalil I, Tantisira K, Weiss S. Systems biology and in vitro validation

identifies family with sequence similarity 129 member A (FAM129A) as an asthma

steroid response modulator. J Allergy Clin Immunol. 2018

Nov;142(5):1479-1488.e12. doi: 10.1016/j.jaci.2017.11.059. Epub 2018 Mar 2.

PubMed PMID: 29410046; PubMed Central PMCID: PMC6111005.

5: Qaisiya M, Mardešić P, Pastore B, Tiribelli C, Bellarosa C. The activation of

autophagy protects neurons and astrocytes against bilirubin-induced cytotoxicity.

Neurosci Lett. 2017 Nov 20;661:96-103. doi: 10.1016/j.neulet.2017.09.056. Epub

2017 Sep 28. PubMed PMID: 28965934.

6: Luo W, Feldman D, McCallister R, Brophy C, Cheung-Flynn J. P2X7R antagonism

after subfailure overstretch injury of blood vessels reverses vasomotor

dysfunction and prevents apoptosis. Purinergic Signal. 2017 Dec;13(4):579-590.

doi: 10.1007/s11302-017-9585-0. Epub 2017 Sep 13. PubMed PMID: 28905300; PubMed

Central PMCID: PMC5714848.

7: Thomas BC, Kay JD, Menon S, Vowler SL, Dawson SN, Bucklow LJ, Luxton HJ,

Johnston T, Massie CE, Pugh M, Warren AY, Barker P, Burling K, Lynch AG, George

A, Burge J, Corcoran M, Stearn S, Lamb AD, Sharma NL, Shaw GL, Neal DE, Whitaker

HC. Whole blood mRNA in prostate cancer reveals a four-gene androgen regulated

panel. Endocr Relat Cancer. 2016 Oct;23(10):797-812. doi: 10.1530/ERC-16-0287.

Epub 2016 Aug 30. PubMed PMID: 27578825.

8: Mendes TB, Nozima BH, Budu A, de Souza RB, Braga Catroxo MH, Delcelo R,

Gazarini ML, Cerutti JM. PVALB diminishes [Ca2+] and alters mitochondrial

features in follicular thyroid carcinoma cells through AKT/GSK3β pathway. Endocr

Relat Cancer. 2016 Sep;23(9):769-82. doi: 10.1530/ERC-16-0181. Epub 2016 Jul 25.

PubMed PMID: 27458244.

9: Shaw GL, Whitaker H, Corcoran M, Dunning MJ, Luxton H, Kay J, Massie CE,

Miller JL, Lamb AD, Ross-Adams H, Russell R, Nelson AW, Eldridge MD, Lynch AG,

Ramos-Montoya A, Mills IG, Taylor AE, Arlt W, Shah N, Warren AY, Neal DE. The

Early Effects of Rapid Androgen Deprivation on Human Prostate Cancer. Eur Urol.

2016 Aug;70(2):214-8. doi: 10.1016/j.eururo.2015.10.042. Epub 2015 Nov 10. PubMed

PMID: 26572708; PubMed Central PMCID: PMC4926724.

10: Carvalheira G, Nozima BH, Cerutti JM. microRNA-106b-mediated down-regulation

of C1orf24 expression induces apoptosis and suppresses invasion of thyroid

cancer. Oncotarget. 2015 Sep 29;6(29):28357-70. doi: 10.18632/oncotarget.4947.

PubMed PMID: 26317551; PubMed Central PMCID: PMC4695065.

11: Yuki R, Aoyama K, Kubota S, Yamaguchi N, Kubota S, Hasegawa H, Morii M, Huang

X, Liu K, Williams R, Fukuda MN, Yamaguchi N. Overexpression of zinc-finger

protein 777 (ZNF777) inhibits proliferation at low cell density through

down-regulation of FAM129A. J Cell Biochem. 2015 Jun;116(6):954-68. doi:

10.1002/jcb.25046. PubMed PMID: 25560148.

12: Koehler JA, Baggio LL, Cao X, Abdulla T, Campbell JE, Secher T, Jelsing J,

Larsen B, Drucker DJ. Glucagon-like peptide-1 receptor agonists increase

pancreatic mass by induction of protein synthesis. Diabetes. 2015

Mar;64(3):1046-56. doi: 10.2337/db14-0883. Epub 2014 Oct 2. PubMed PMID:

25277394.

13: Yick CY, Zwinderman AH, Kunst PW, Grünberg K, Mauad T, Chowdhury S, Bel EH,

Baas F, Lutter R, Sterk PJ. Gene expression profiling of laser microdissected

airway smooth muscle tissue in asthma and atopy. Allergy. 2014 Sep;69(9):1233-40.

doi: 10.1111/all.12452. Epub 2014 Jul 19. PubMed PMID: 24888725.

14: Liu J, Qin J, Mei W, Zhang H, Yuan Q, Peng Z, Luo R, Yuan X, Huang L, Tao L.

Expression of Niban in renal interstitial fibrosis. Nephrology (Carlton). 2014

Aug;19(8):479-89. doi: 10.1111/nep.12266. PubMed PMID: 24750539.

15: Sigstad E, Paus E, Bjøro T, Berner A, Grøholt KK, Jørgensen LH,

Sobrinho-Simões M, Holm R, Warren DJ. Reply to 'The new molecular markers DDIT3,

STT3A, ARG2 and FAM129A are not useful in diagnosing thyroid follicular tumors'.

Mod Pathol. 2013 Apr;26(4):613-5. doi: 10.1038/modpathol.2013.39. PubMed PMID:

23542526.

16: Carvalheira GM, Nozima BH, Riggins GJ, Cerutti JM. DDIT3, STT3A (ITM1), ARG2

and FAM129A (Niban, C1orf24) in diagnosing thyroid carcinoma: variables that may

affect the performance of this antibody-based test and promise. Mod Pathol. 2013

Apr;26(4):611-3. doi: 10.1038/modpathol.2012.212. PubMed PMID: 23542525.

17: Yick CY, Zwinderman AH, Kunst PW, Grünberg K, Mauad T, Fluiter K, Bel EH,

Lutter R, Baas F, Sterk PJ. Glucocorticoid-induced changes in gene expression of

airway smooth muscle in patients with asthma. Am J Respir Crit Care Med. 2013 May

15;187(10):1076-84. doi: 10.1164/rccm.201210-1886OC. PubMed PMID: 23491407.

18: Cerutti JM. Employing genetic markers to improve diagnosis of thyroid tumor

fine needle biopsy. Curr Genomics. 2011 Dec;12(8):589-96. doi:

10.2174/138920211798120781. PubMed PMID: 22654558; PubMed Central PMCID:

PMC3271311.

19: Ji H, Ding Z, Hawke D, Xing D, Jiang BH, Mills GB, Lu Z. AKT-dependent

phosphorylation of Niban regulates nucleophosmin- and MDM2-mediated p53 stability

and cell apoptosis. EMBO Rep. 2012 Jun 1;13(6):554-60. doi:

10.1038/embor.2012.53. Erratum in: EMBO Rep. 2014 Sep;15(9):1000. PubMed PMID:

22510990; PubMed Central PMCID: PMC3367238.

20: Sigstad E, Paus E, Bjøro T, Berner A, Grøholt KK, Jørgensen LH,

Sobrinho-Simões M, Holm R, Warren DJ. The new molecular markers DDIT3, STT3A,

ARG2 and FAM129A are not useful in diagnosing thyroid follicular tumors. Mod

Pathol. 2012 Apr;25(4):537-47. doi: 10.1038/modpathol.2011.188. Epub 2011 Dec 9.

PubMed PMID: 22157935; PubMed Central PMCID: PMC3318159.

21: Miller S, Rogers HA, Lyon P, Rand V, Adamowicz-Brice M, Clifford SC, Hayden

JT, Dyer S, Pfister S, Korshunov A, Brundler MA, Lowe J, Coyle B, Grundy RG.

Genome-wide molecular characterization of central nervous system primitive

neuroectodermal tumor and pineoblastoma. Neuro Oncol. 2011 Aug;13(8):866-79. doi:

10.1093/neuonc/nor070. PubMed PMID: 21798848; PubMed Central PMCID: PMC3145471.

22: Patel MR, Stadler ME, Deal AM, Kim HS, Shores CG, Zanation AM. STT3A,

C1orf24, TFF3: putative markers for characterization of follicular thyroid

neoplasms from fine-needle aspirates. Laryngoscope. 2011 May;121(5):983-9. doi:

10.1002/lary.21736. PubMed PMID: 21520112.

23: Cerutti JM, Oler G, Delcelo R, Gerardt R, Michaluart P Jr, de Souza SJ,

Galante PA, Huang P, Riggins GJ. PVALB, a new Hürthle adenoma diagnostic marker

identified through gene expression. J Clin Endocrinol Metab. 2011

Jan;96(1):E151-60. doi: 10.1210/jc.2010-1318. Epub 2010 Oct 6. PubMed PMID:

20926528; PubMed Central PMCID: PMC3038489.

24: Ito S, Fujii H, Matsumoto T, Abe M, Ikeda K, Hino O. Frequent expression of

Niban in head and neck squamous cell carcinoma and squamous dysplasia. Head Neck.

2010 Jan;32(1):96-103. doi: 10.1002/hed.21153. PubMed PMID: 19536772.

25: Sun GD, Kobayashi T, Abe M, Tada N, Adachi H, Shiota A, Totsuka Y, Hino O.

The endoplasmic reticulum stress-inducible protein Niban regulates eIF2alpha and

S6K1/4E-BP1 phosphorylation. Biochem Biophys Res Commun. 2007 Aug

17;360(1):181-7. Epub 2007 Jun 12. PubMed PMID: 17588536.

26: Matsumoto F, Fujii H, Abe M, Kajino K, Kobayashi T, Matsumoto T, Ikeda K,

Hino O. A novel tumor marker, Niban, is expressed in subsets of thyroid tumors

and Hashimoto's thyroiditis. Hum Pathol. 2006 Dec;37(12):1592-600. Epub 2006 Sep

1. PubMed PMID: 16949643.

27: Cerutti JM, Latini FR, Nakabashi C, Delcelo R, Andrade VP, Amadei MJ, Maciel

RM, Hojaij FC, Hollis D, Shoemaker J, Riggins GJ. Diagnosis of suspicious thyroid

nodules using four protein biomarkers. Clin Cancer Res. 2006 Jun 1;12(11 Pt

1):3311-8. PubMed PMID: 16740752.

28: Maciel RM, Kimura ET, Cerutti JM. [Pathogenesis of differentiated thyroid

cancer (papillary and follicular)]. Arq Bras Endocrinol Metabol. 2005

Oct;49(5):691-700. Epub 2006 Jan 23. Review. Portuguese. PubMed PMID: 16444351.

29: Kannangai R, Diehl AM, Sicklick J, Rojkind M, Thomas D, Torbenson M. Hepatic

angiomyolipoma and hepatic stellate cells share a similar gene expression

profile. Hum Pathol. 2005 Apr;36(4):341-7. PubMed PMID: 15891994.

30: Hino O. Multistep renal carcinogenesis in the Eker (Tsc 2 gene mutant) rat

model. Curr Mol Med. 2004 Dec;4(8):807-11. Review. PubMed PMID: 15579027.

31: Cerutti JM, Delcelo R, Amadei MJ, Nakabashi C, Maciel RM, Peterson B,

Shoemaker J, Riggins GJ. A preoperative diagnostic test that distinguishes benign

from malignant thyroid carcinoma based on gene expression. J Clin Invest. 2004

Apr;113(8):1234-42. PubMed PMID: 15085203; PubMed Central PMCID: PMC385398.

32: Adachi H, Majima S, Kon S, Kobayashi T, Kajino K, Mitani H, Hirayama Y,

Shiina H, Igawa M, Hino O. Niban gene is commonly expressed in the renal tumors:

a new candidate marker for renal carcinogenesis. Oncogene. 2004 Apr

22;23(19):3495-500. PubMed PMID: 14990989.

33: Hooper-Bùi LM, Appel AG, Rust MK. Preference of food particle size among

several urban ant species. J Econ Entomol. 2002 Dec;95(6):1222-8. PubMed PMID:

12539835.

34: Tripp JM, Suiter DR, Bennett GW, Klotz JH, Reid BL. Evaluation of control

measures for black carpenter ant (Hymenoptera: Formicidae). J Econ Entomol. 2000

Oct;93(5):1493-7. PubMed PMID: 11057723.

35: Majima S, Kajino K, Fukuda T, Otsuka F, Hino O. A novel gene "Niban"

upregulated in renal carcinogenesis: cloning by the cDNA-amplified fragment

length polymorphism approach. Jpn J Cancer Res. 2000 Sep;91(9):869-74. PubMed

PMID: 11011112; PubMed Central PMCID: PMC5926447.
